# HalluDesign-NA: Extending HalluDesign for De Novo Nucleic Acid Design

**DOI:** 10.64898/2026.06.10.730767

**Authors:** Minchao Fang, Zhe Wang, Longxing Cao

**Affiliations:** State Key Laboratory of Gene Expression, School of Life Sciences, Westlake University, Hangzhou, China; Artificial Intelligence Drug Design Core Laboratory, Westlake Laboratory of Life Sciences and Biomedicine, Hangzhou, China

## Abstract

AlphaFold3 has revolutionized the prediction of biomolecular structures and interactions, including atomic-level modeling of nucleic acids. However, the de novo design of structured and functional nucleic acids remains a significant challenge. Here, we extend our HalluDesign framework to nucleic acid design by integrating NA-MPNN for nucleic acid sequence optimization and design. This new framework, HalluDesign-NA, enables iterative sequence-structure co-optimization, facilitating the de novo design of nucleic acids. Computational benchmarking across ssDNA, ssRNA, and aptamer design tasks demonstrates consistent improvements in confidence scores (pLDDT, ipTM), supporting the feasibility of de novo nucleic acid design under various constraints, such as sequence length, symmetry, and protein structure context. We anticipate that HalluDesign-NA will accelerate the de novo design of functional nucleic acids for applications in biotechnology and medicine. The source code for HalluDesign-NA is available at https://github.com/MinchaoFang/HalluDesign_NA.

## Background

Deep learning–based foundation models have dramatically advanced protein structure prediction, enabling atomic-level modeling from sequence alone (*1-3*). Recent models, such as AlphaFold3 (*4*) and its open-source variants (*5*), have further expanded this capability beyond proteins, allowing more accurate prediction of nucleic acid structures and protein–nucleic acid complexes than before (*6, 7*). Alongside these developments, generative models like RFpoly (*8*) and Odesign (*9*) have emerged to enable nucleic acid design. However, the de novo design of functional nucleic aptamers capable of interacting with other molecules using these generative models remains challenging, and, to date, no experimental validation has been reported. While several promising sequence-based methods have been developed for generating functional aptamers, they typically rely on high-throughput experiments (*10, 11*). Currently, functional nucleic acids are primarily obtained through high-throughput screening approaches such as SELEX. Nevertheless, researchers are increasingly exploring the potential for de novo functional nucleic acid design in low-throughput settings. Despite these efforts, the de novo design of functional nucleic acids remains far behind that of functional proteins, which can now be designed to target a diverse range of functionalities.

HalluDesign is a general framework for protein optimization and de novo design, built upon the inherent “hallucination” effect of the generative diffusion module in AF3-style models (*12*). By integrating a sequence design module, such as LigandMPNN or ProteinMPNN (*13, 14*), with a structure prediction model like AlphaFold3, HalluDesign enables iterative sequence-structure co-optimization. We demonstrated the broad applicability and high efficiency of HalluDesign in generating new protein structures, designing protein binders targeting various biomolecules including ssRNA, and developing enzymes. Building on HalluDesign’s strong performance in protein optimization and design, we sought to investigate whether AlphaFold3’s high precision could facilitate high-quality design of functional nucleic acids. With the development of NA-MPNN (*15*)—a model based on the MPNN framework that enables nucleic acid sequence design—we explored whether NA-MPNN could replace the protein sequence design ProteinMPNN/LigandMPNN module in HalluDesign, thereby extending its capabilities from protein to functional nucleic acid design.

### HalluDesign-NA Framework

HalluDesign-NA employs NA-MPNN (*15*) to update nucleic acid sequences, replacing the original protein sequence design model (ProteinMPNN or LigandMPNN (*13, 14*)). The rest of the framework remains unchanged (**Fig. 1**). For more details, please refer to our HalluDesign work (*12*). We develop HalluDesign-NA implementions on both the foundation model Protenix and AF3. And during the design process, we use random sequence initialization to generate the starting sequences for both DNA and RNA design.

**Fig. 1.**
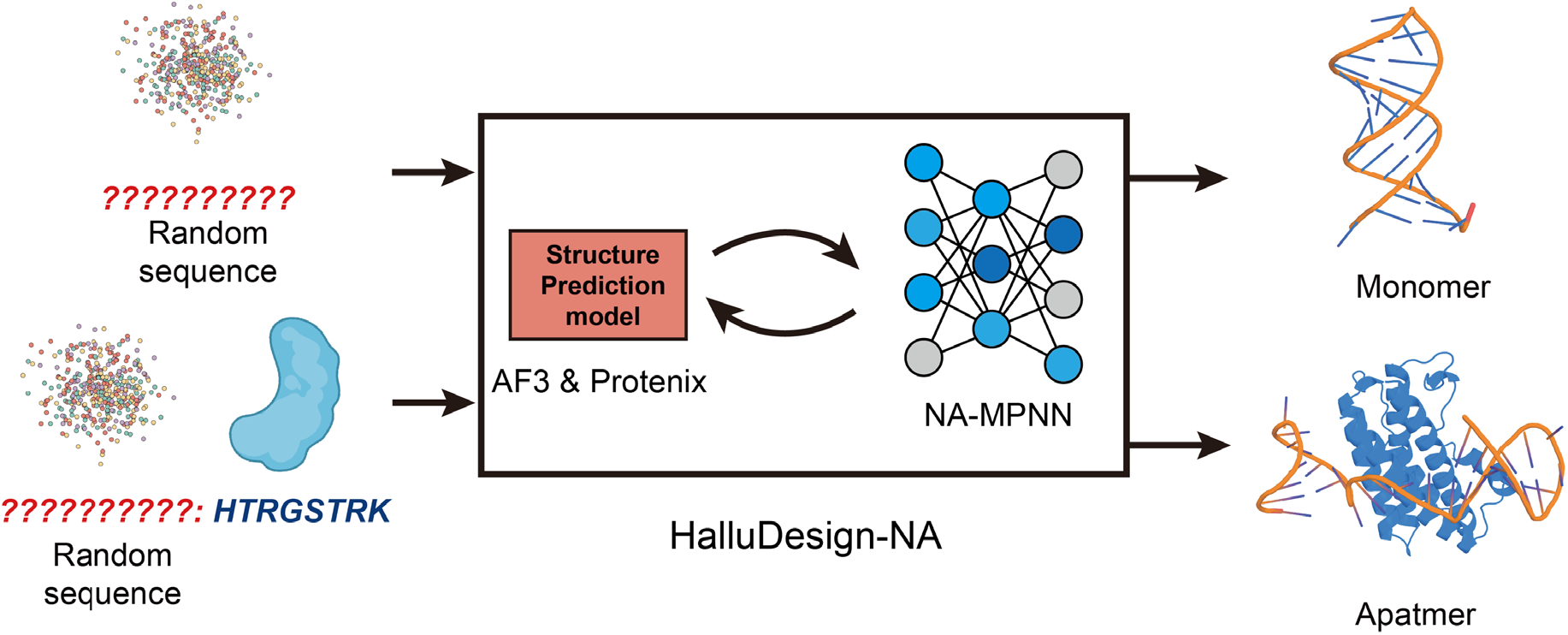
Overview of the HalluDesign-NA framework. Compared with the original HalluDesign framework, the MPNN protein sequence design module is replaced by NA-MPNN, a nucleic acid sequence design module. During aptamer design, the protein target is kept fixed.

### Computational Design of DNA and RNA structures

We first used HalluDesign-NA to design single-stranded DNA (ssDNA) and RNA (ssRNA) monomers, exploring sequence lengths ranging from 50 to 80 nucleotides (**Fig. 2**). Starting from randomly initialized sequences, HalluDesign-NA consistently improved computational metrics of the nucleic acid structures, including pLDDT and pTM. The final designs displayed a variety of base pairing patterns and compact structures (**Fig. 3A**), demonstrating the feasibility of HalluDesign-NA for nucleic acid design. This approach supports the design of nucleic acid monomers with lengths exceeding 300 nucleotides, with some monomers achieving pLDDT scores of up to 80 (**Fig. 3B**). For symmetric design, by incorporating NA-MPNN symmetry sequence constraints during sequence design, HalluDesign-NA is able to generate symmetric structures. For example, the symmetric dimer designs achieve a pLDDT score of up to 80 (**Fig. 3C**), indicating a high level of structural confidence.

**Fig. 2.**
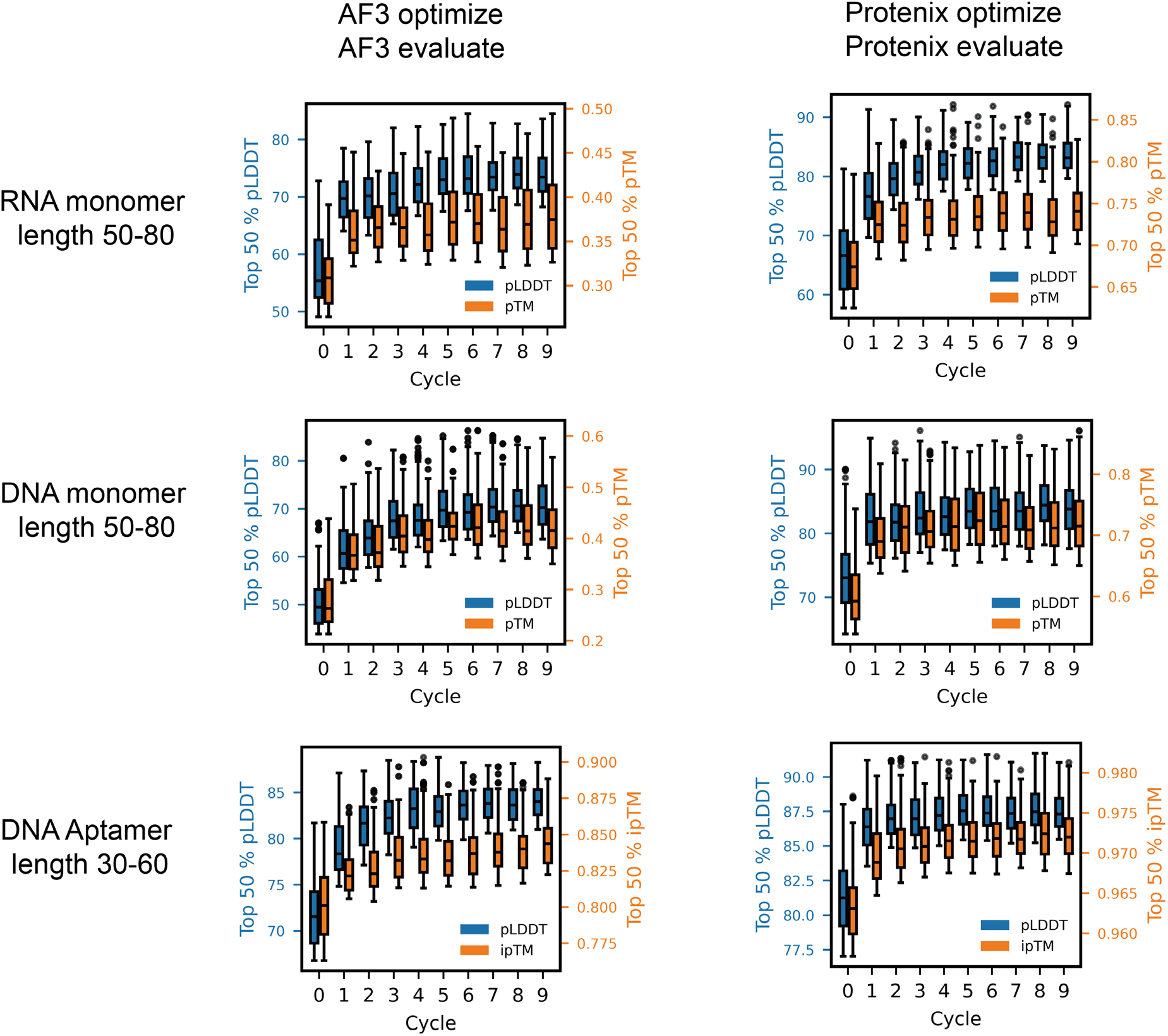
HalluDesign-NA benchmark on ssDNA monomer and ssRNA and ssDNA aptamer from scratch design using Protenix or AF3 model.

**Fig. 3.**
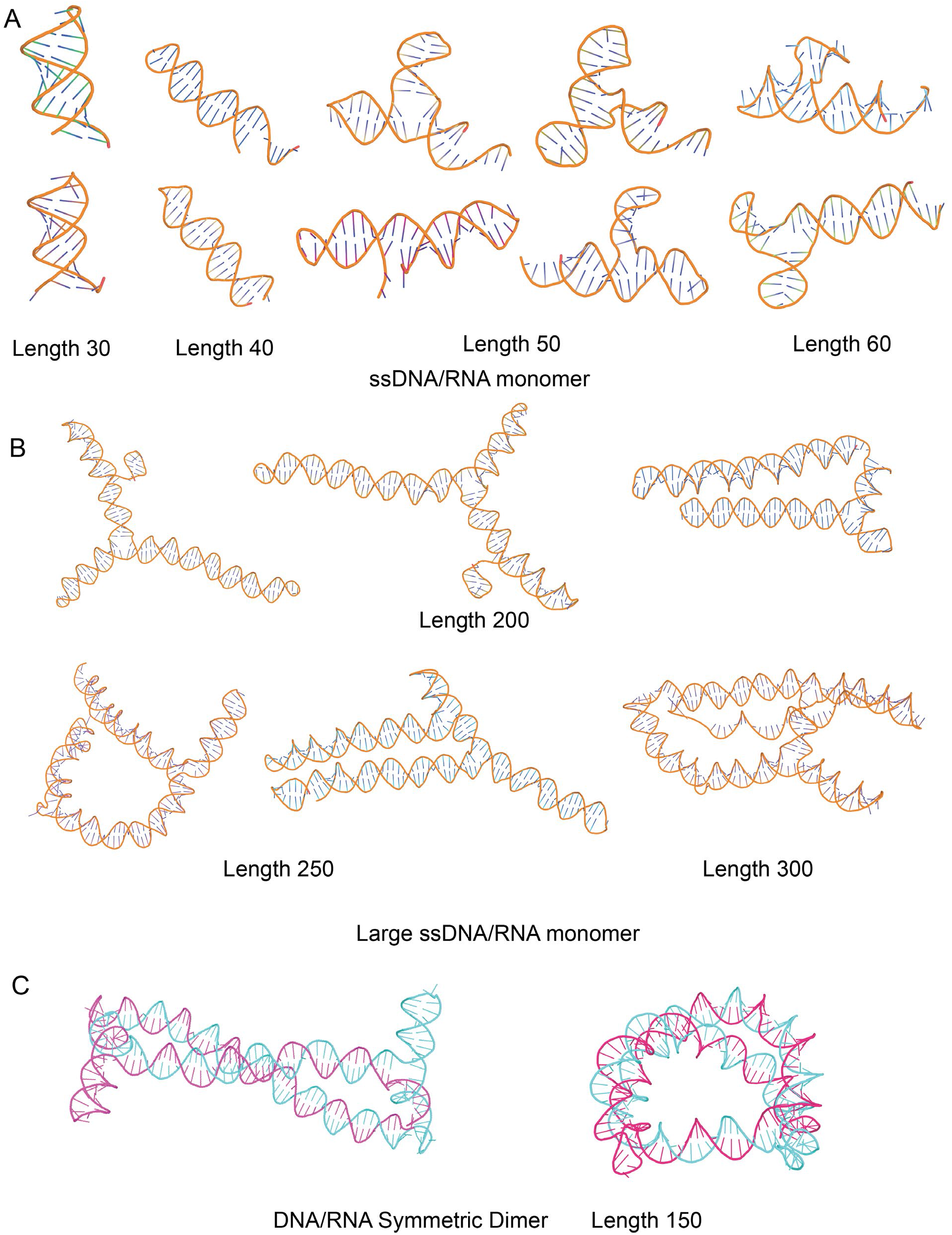
HalluDesign-NA generates diverse topologies of ssRNA, ssDNA, and symmetric DNA dimers. (**A**) Designs of short ssDNA and ssRNA with lengths of 30-60 nucleotides. (**B**) Designs of long ssDNA and ssRNA with lengths of 200-300 nucleotides. (**C**) Symmetric DNA dimer design with a length of 150 nucleotides.

### Computational Design of protein binding DNA aptamers

We next applied HalluDesign-NA to design DNA aptamers targeting IL6, exploring aptamer lengths ranging from 30 to 60 nucleotides (**Fig. 2**). Similarly, the AF3-style model Protenix and AF3 both can be served as the foundational structure prediction model, and nucleic acid sequences were randomly initialized to generate the initial design models. HalluDesign-NA consistently optimized computational metrics over iterative design cycles, with the protein sequence held fixed in ssDNA aptamer design (**Fig. 4**). Notably, the designed binding site mostly overlaps with the aptamer binding site identified in the crystal structure (PDB ID: 4NI7). We are exploring optimal computational metrics for selecting designs for experimental characterization, as well as experimentally characterizing the designed aptamers, with the goal of advancing the structural-level design of functional nucleic acids.

**Fig. 4.**
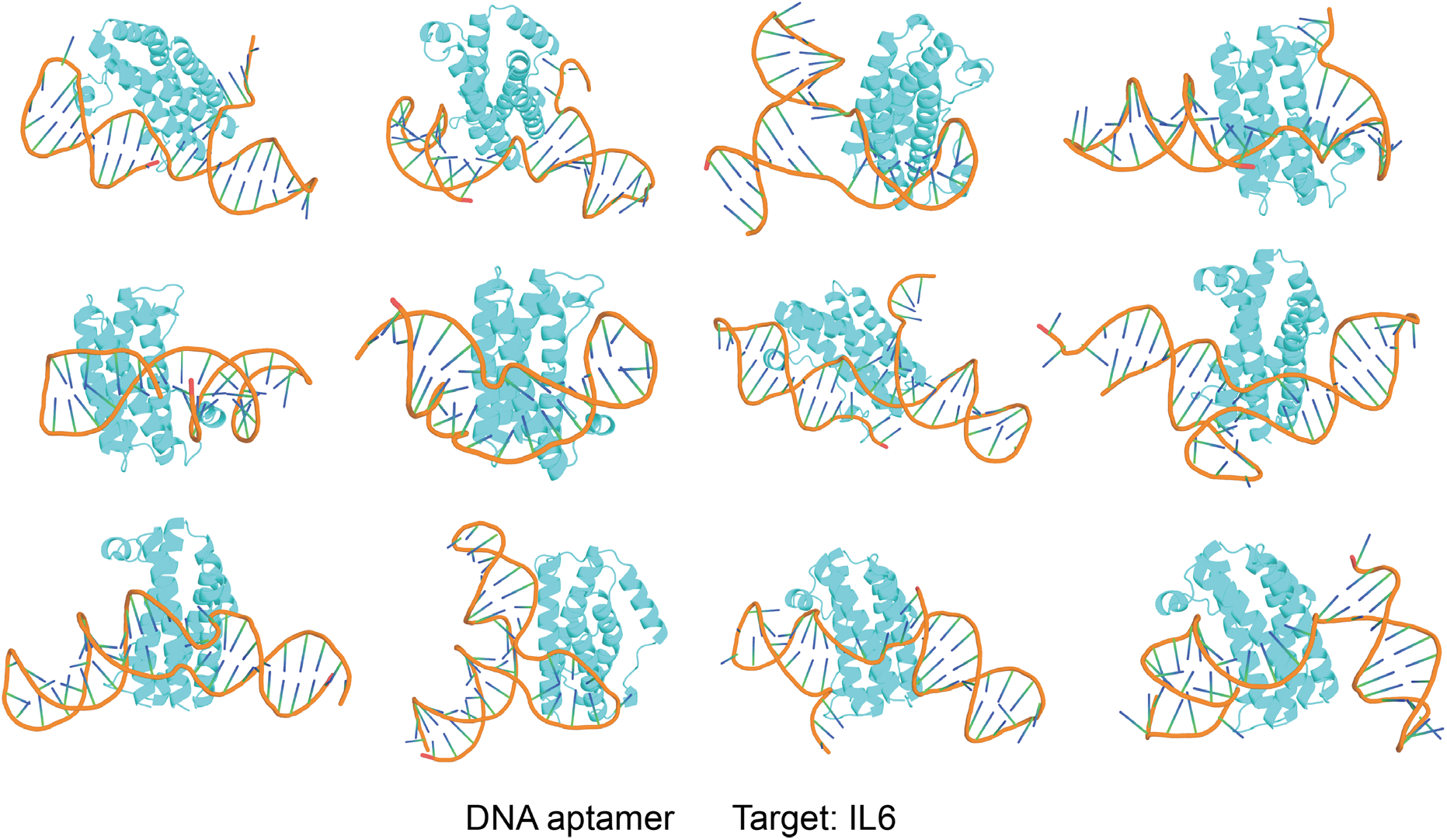
HalluDesign-NA generates diverse topologies of IL-6-targeting aptamers with lengths of 30-60 nucleotides. The IL-6 protein target is shown in gray and green, while the aptamer is colored orange.

## Discussion

De novo design of functional nucleic acids or aptamers in low-throughput scenarios remains far from routine. HalluDesign-NA offers a new approach for designing functional nucleic acids, leveraging state-of-the-art structure prediction models. Although our in silico benchmark demonstrates its potential, several obstacles must still be overcome to ensure experimental success. One major challenge is the limited availability of resolved nucleic acid–protein complexes, or even isolated nucleic acid structures, especially when compared to the vast number of protein structures available in the PDB (*16*). While current aptamer selection methods support efficient, high-throughput sequence identification, the majority of experimental results remain sequence-based (*17*). Sequence models can learn sequence-level representations, yet reliably transferring these features to the structural domain—particularly for design—remains difficult.

Moreover, the secondary and tertiary structures of nucleic acids are highly sensitive to environmental factors such as temperature, buffer composition, and ionic strength (*18, 19*). Variations in cooling rate (rapid versus slow) and subtle changes in solution conditions can significantly alter experimental structural outcomes, which may diverge from those found in physiological environments. Unfortunately, many existing datasets do not provide detailed documentation of these experimental parameters, further complicating the reproducibility and reliability of structure prediction and de novo design—especially in the context of limited training data.

## Conclusion

HalluDesign has proven to be a versatile and extensible framework, with demonstrated applications ranging from protein optimization to de novo enzyme generation. HalluDesign-NA further expands the framework’s capabilities to nucleic acid design. Our in silico benchmark results show that HalluDesign-NA can generate nucleic acids with high-confidence metrics, providing researchers with promising starting points for experimental validation. As HalluDesign-NA extends generative design to nucleic acids, we anticipate broad applications in molecular diagnostics, drug development, and synthetic biotechnology.

## Code availability

All source code developed in this study is available at Github website. HalluDesign-NA (https://github.com/MinchaoFang/HalluDesign_NA). HalluDesign (https://github.com/MinchaoFang/HalluDesign).

